# Human-scATAC-Corpus: a comprehensive database of scATAC-seq data

**DOI:** 10.1101/2025.09.05.674505

**Authors:** Xiaoyang Chen, Zijing Gao, Keyi Li, Zian Wang, Qun Jiang, Xuejian Cui, Zhen Li, Rui Jiang

**Author notes:** Joint First Authors (Xiaoyang Chen, Zijing Gao, Keyi Li, Zian Wang).

## Abstract

Single-cell Assay for Transposase-Accessible Chromatin using sequencing (scATAC-seq) profiles chromatin accessibility at cellular resolution, making it possible to reveal epigenomic landscapes that govern gene regulation in a variety of cells. Nevertheless, heterogeneous feature spaces and complex processing pipelines have impeded the construction of an ensemble resource capable of supporting diverse downstream analytical scenarios. To address this gap, we present Human-scATAC-Corpus (https://health.tsinghua.edu.cn/human-scatac-corpus/), a comprehensive database of human scATAC-seq comprising 5,407,621 cells from 35 datasets across 37 tissues or cell lines. To support complementary use cases, each dataset is distributed in three aligned formats: cell-by-candidate cis-regulatory element matrices for cross-dataset integration, raw fragment files for flexible processing, and cell-by-peak matrices for within-dataset analyses. This resource spans diverse biological contexts and includes rich metadata, enabling method benchmarking and development, as well as pretraining of foundation models. The website offers searchable browsing, detailed dataset pages, on-demand downloads, and tutorials. EpiAgent, a foundation model pretrained on Human-scATAC-Corpus, is further integrated to provide online analyses, including reference mapping, embedding extraction, and cell type annotation. Human-scATAC-Corpus establishes a unified and scalable substrate for single-cell epigenomics and is intended to accelerate discovery while standardizing evaluation across tasks.

**GRAPHICAL ABSTRACT:** 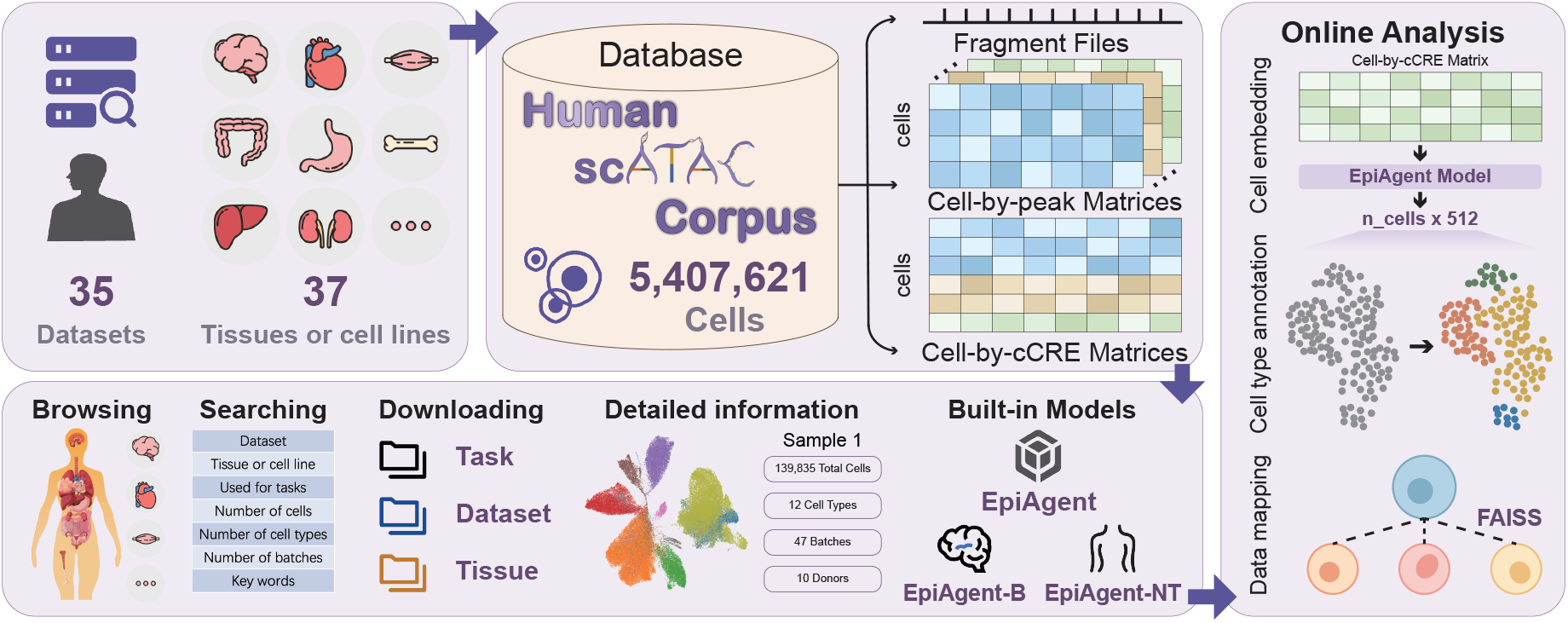

## INTRODUCTION

Single-cell Assay for Transposase-Accessible Chromatin using sequencing (scATAC-seq) enables the characterization of chromatin accessibility at the resolution of individual cells, thereby revealing the regulatory landscape at the cellular level (1). In a broad range of biomedical researches, including investigations of cellular heterogeneity (2), tissue differentiation (3), and disease pathogenesis (4), scATAC-seq data have been serving as an essential and irreplaceable resource. Recent advances in sequencing technologies have driven two major developments in this field. First, improvements in throughput have led to the development of high-throughput techniques such as sci-ATAC-seq (5) and sci-ATAC-seq3 (6), enabling the construction of large-scale cell atlases spanning fetal development (6), adult tissues (7), and the adult brain (8). Second, the scope of applications has been expanding, with techniques such as CRISPR-sciATAC (9) and Spear-ATAC (10) integrating CRISPR-based genome editing to capture cell state changes under various perturbations. Indeed, the growth in both data scale and application diversity has fueled the emergence of task-specific analytical methods and the development of foundation models trained on scATAC-seq data. However, these efforts, no matter focusing on cell-level foundation models such as EpiAgent (11), concentrating on genome-level foundation models such as GET (12), or aiming at developing task-specific models (13-20), all require a publicly accessible, ensemble database that encompasses the diversity of scATAC-seq applications and tissue types.

In the single-cell field, large-scale ensemble databases already exist for other omics. For example, the scRNA-seq–based Cell x Gene repository (21) has driven the adoption of large-scale pretraining paradigms for transcriptomic data (22-27). scRNA-seq features are anchored on a shared gene set, and thus each dataset can be represented as a cell-by-gene matrix that supports most analytic workflows. In contrast, scATAC-seq lacks a universal feature space. Data are typically stored as raw fragment files or as cell-by-peak matrices whose peak definitions differ across datasets (28). Consequently, no single storage format can satisfy all analytical requirements, and multiple data representations are often necessary. Existing scATAC-seq ensemble resources, such as scATAC-Ref (29), have reduced the time required for users to locate data and facilitated the development of related methods, yet they still present several notable limitations. First, reliance on cell-by-peak matrices yields incompatible feature spaces that impede joint analysis. Second, current scale is insufficient for modern pretraining regimes. Third, data modalities and metadata are narrow, restricting the breadth and fidelity of downstream benchmarking. Fourth, online analysis capabilities are constrained, with few powerful embedded models or reusable applications applied to the hosted data. These limitations underscore the urgent need for a comprehensive database of scATAC-seq data that can address the above challenges.

To facilitate research in single-cell epigenomics, we developed Human-scATAC-Corpus (https://health.tsinghua.edu.cn/human-scatac-corpus/), a comprehensive database of scATAC-seq data. The current release contains 5,407,621 cells from 35 datasets across 37 human tissues or cell lines. All data are provided in three complementary formats: cell-by-candidate cis-regulatory element (cCRE) matrices for integrative multi-dataset analysis, fragment files for flexible user-defined processing, and cell-by-peak matrices for within-dataset analysis. The collection spans diverse biological contexts, enabling the use in a wide range of downstream tasks, from method development and benchmarking to large-scale pretraining of foundation models. Human-scATAC-Corpus also integrates EpiAgent, a foundation model pretrained on this resource, supporting online analyses including data mapping to reference, cell embedding extraction, and cell type annotation via supervised derivatives EpiAgent-B and EpiAgent-NT. Other functionalities include user-friendly and intuitive browsing, structured and keyword-based search, in-depth dataset detail, on-demand download, customized tutorials, and interactive online analysis. We anticipate that Human-scATAC-Corpus will provide a versatile and scalable resource to accelerate discoveries in single-cell epigenomics and advance the development of analytical methods.

## DATA COLLECTION AND DATABASE CONTENT

### Data collection

For the collection of scATAC-seq data, we implemented a two-stage workflow consisting of literature and repository searches followed by eligibility screening and preliminary quality control (Figure 1). First, we queried the PubMed database and the Gene Expression Omnibus (GEO) repository using the keywords “ Human” and “ scATAC-seq”, yielding a total of 205 published studies. For these candidate studies, we examined whether the datasets contained scATAC-seq data derived from healthy human samples and excluded datasets that did not meet this criterion. Because the primary objective of our database is to provide high-quality pretraining resources for the foundation model, we excluded cancer and other pathological conditions, as such cells may display unknown regulatory states that could introduce noise and reduce the reliability of training on normal cells.

**Figure 1.**
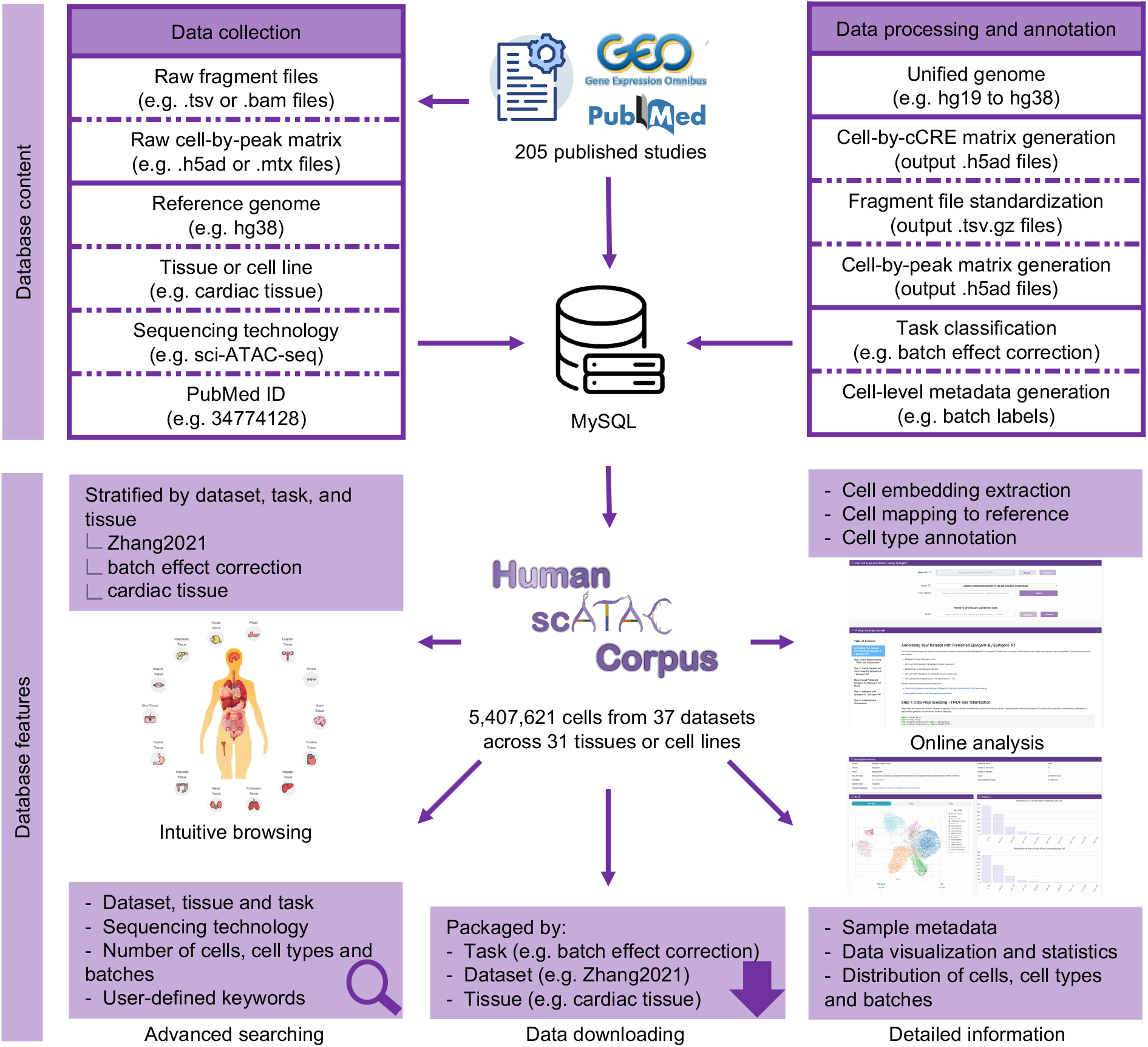
Overview of data collection, data processing and annotation, and database features of Human-scATAC-Corpus. Cell type annotation

At the same time, we retained widely used benchmark datasets, such as Buenrostro2018 (30), as well as other cells reported in the studies that contain normal cells, to ensure that our resource remains suitable for both method development and benchmarking. For the studies retained after screening, we downloaded the publicly available data and further examined whether fragment files or high-quality cell-by-peak matrices were provided. Ultimately, we curated and collected 35 datasets spanning 37 tissues or cell lines. For each dataset, we additionally recorded essential metadata, including the reference genome, tissue or cell line of origin, sequencing technology, and PubMed identifier (PMID).

### Data processing and annotation

For the Human-scATAC-Corpus, only data aligned to the reference genome GRCh38/hg38 are currently provided. Consequently, the first step involves mapping data originally with GRCh37/hg19 into GRCh38/hg38 by liftOver (31). To facilitate usability and ensure accurate data retrieval, we defined the minimal storage unit (also denoted as a sample) in Human-scATAC-Corpus as a single tissue within a dataset. Each sample is named in the format dataset– tissue/cell line and assigned a unique identifier (ID) for reference. At the sample level, we also annotated their detailed information based on the available metadata.

For each sample, Human-scATAC-Corpus offers three data formats: a cell-by-cCRE matrix, fragment files, and a cell-by-peak matrix. The cell-by-cCRE matrix is provided for all samples in Human-scATAC-Corpus, while the other two depend on the availability of source data. Fragment files, typically stored in formats such as.tsv or.bam, and cell-by-peak matrices, commonly stored in.h5ad or.mtx formats, are the most frequently used data types in current scATAC-seq studies. During the processing stage, we standardized all fragment files into a series of compressed.tsv.gz files, with each file corresponding to a batch. Cell-by-peak matrices were processed into a single.h5ad file. Since fragment files represent relatively non-processed data and cell-by-peak matrices are dataset-specific due to non-unified peak definitions, they are generally less suitable for integrative analysis across datasets. To address this, we constructed a unified set of cCREs by taking the union of cCREs or peak lists from three large-scale datasets spanning distinct tissues (7,8,32). Fragment files (or, when unavailable, a cell-by-peak matrix) were then mapped to this unified cCRE set to generate a cell-by-cCRE matrix of.h5ad format, consistent with the strategy employed in our EpiAgent study (11). In that work, we demonstrated that these cCREs exhibit substantial overlap with peaks across the majority of datasets, and the superior performance of EpiAgent across multiple datasets and tasks further supports the reliability of the cCRE definition. After mapping, we applied quality filtering by removing cells with fewer than 100 nonzero values in the cell-by-cCRE matrix, and we ensured consistency of cell storage across all three data formats. Sample names and batch information were incorporated into file and barcode prefixes, and standardized naming conventions were applied to facilitate downstream use. Given that scATAC-seq data formats vary considerably across studies, these steps are both labor-intensive and dataset-specific. Standardization and enhanced data usability therefore represent core strengths of Human-scATAC-Corpus.

Finally, we classified samples according to the availability of cell-level metadata to indicate the types of tasks they support. Specifically, we designated samples for (i) self-supervised pretraining when all cells are included, (ii) unsupervised feature extraction and supervised cell type annotation when cell-type labels are available, (iii) batch effect correction when batch labels is provided, (iv) prediction of cell states under unseen genetic perturbation when CRISPR-based perturbation information is available, and (v) prediction of cell states under out-of-sample stimulated perturbation when external stimulation labels are included. To ensure interoperability, all cell-level metadata were standardized: for cell-by-cCRE and cell-by-peak matrices, metadata are stored in the.obs field of the Anndata object (33), while for fragment files, metadata are provided as separate.csv files. This design facilitates the development and benchmarking of analytic methods across diverse tasks (Figure 1).

### Database statistics

As of the current release, the Human-scATAC-Corpus database contains 5,407,621 cells (approximately 5.4 million), organized into 77 samples defined by “ dataset–tissue/cell line” combinations and encompassing 9 sequencing technologies. Among these, brain tissue accounts for the largest proportion, with 2,549,071 cells, representing 47.14% of the total. By data type, the database provides cell-by-cCRE matrices for all 77 samples (covering all cells), fragment files for 68 samples (4,752,227 cells in total), and cell-by-peak matrices for 65 samples (4,117,861 cells in total). The Human-scATAC-Corpus database supports method development and benchmarking for at least five types of analytic tasks: all cells can be used for the self-supervised pretraining task; 66 samples (5,192,122 cells) contain batch labels, enabling the batch effect correction task; 46 samples (3,021,043 cells) have cell type annotations, supporting unsupervised feature extraction and supervised cell type annotation; 4 samples (167,064 cells) are suitable for cell states prediction under unseen genetic perturbation; and another 4 samples (89,723 cells) are suitable for cell states prediction under out-of-sample stimulated perturbation.

### Built-in foundation model

Prior databases have largely focused on data provision for download and offline use, with little or no capacity for online analysis, or only limited functionality based on simple plug-ins. As a result, the information embedded in the data has often been underutilized. In our study, we integrate into the server side of the database a pretrained EpiAgent model, trained on a subset of Human-scATAC-Corpus, together with two supervised derivatives: EpiAgent-B for the annotation of cells from brain tissues and EpiAgent-NT for the annotation of cells from non-brain tissues (11). This integration enables users to directly perform online analyses, thereby enhancing the accessibility and utility of the database.

## DATABASE FEATURES AND APPLICATIONS

### User-friendly browsing

We developed an intuitive web interface that hosts an unprecedented repository of human scATAC-seq data, empowering users to advance research in single-cell epigenomics and to perform online analyses with a built-in foundation model, EpiAgent (11). The Home page features interactive images of human anatomy and comprehensive statistics detailing the distribution of over 5.4 million cells across file formats, tissues, and analytical tasks, alongside related studies and databases pertinent to single-cell epigenomics. By clicking on the anatomical image, users are directed to the *Browse* page, enabling them to explore and download scATAC-seq data associated with the corresponding tissues (Figure. 2A). At the *Browse* page, the datasets are categorized in parallel by three key attributes: tissue or cell line, dataset, and task. As shown in Figure 2B, Human-scATAC-Corpus provides an interactive metadata table, a visualized UMAP projection from cell embeddings generated by EpiAgent, and comprehensive statistics for metadata. The metadata table presents basic information for each sample, including tissue, organ, PMID, hyperlinks to source data and publication, sequencing technology, and raw data type. Users can directly download the cell-by-peak matrix, cell-by-cCRE matrix, and fragment files via clickable icons in the table. Each sample is also assigned an interactive unique ID that directs to a new webpage with detailed sample information. The UMAP visualization displays up to 100,000 sampled cells of the selected item, labelled by cell type, batch, or donor, with interactive label switching. The statistics comprise pie charts illustrating the distribution of cell counts by sequencing technology, raw data type, tissue, and dataset.

**Figure 2.**
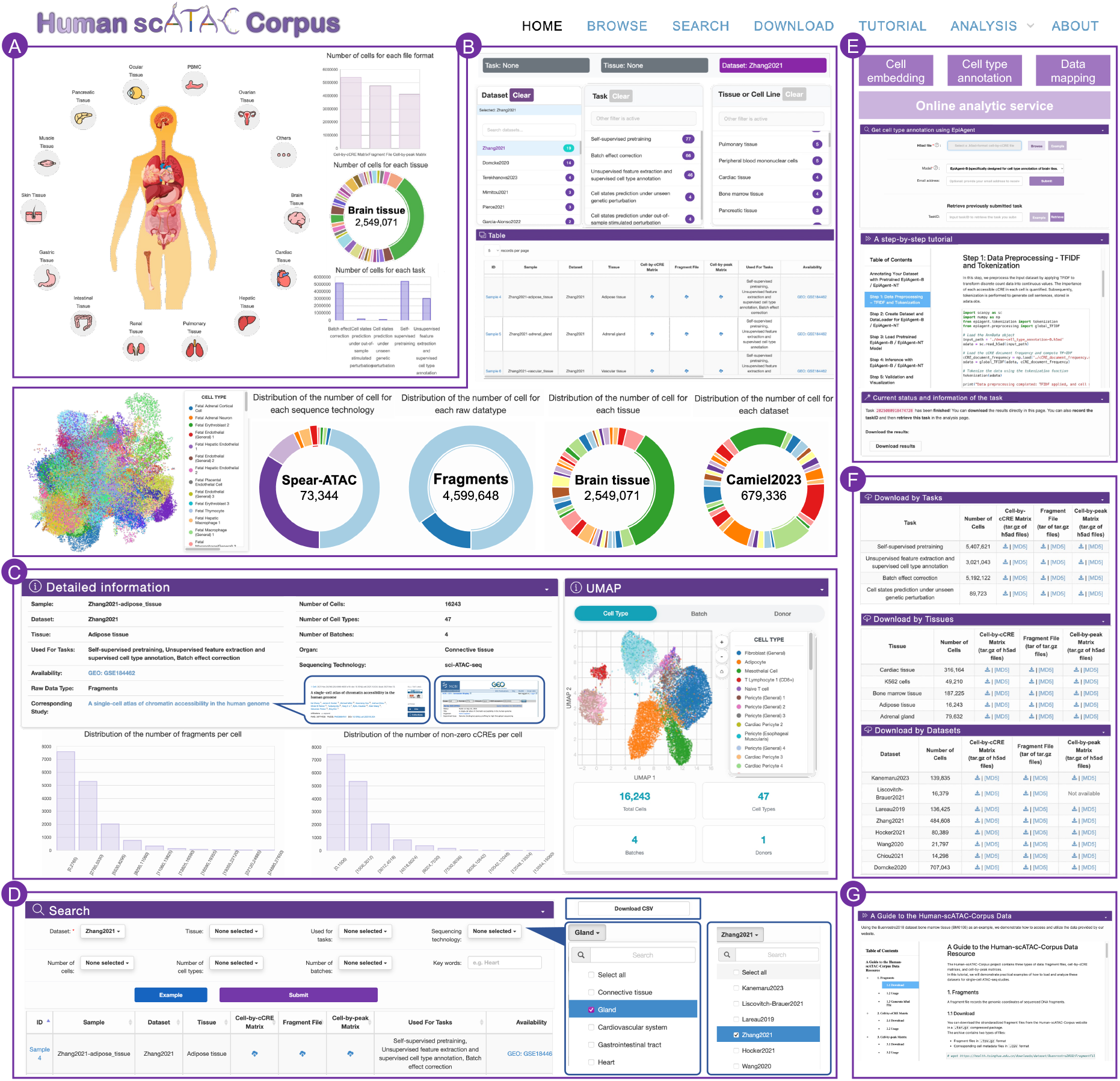
The schematic features on various webpages of Human-scATAC-Corpus. **(A)** *Home*: interactive images of human tissue-level navigation and overview data statistics. **(B)** *Browse*: hierarchical organization of datasets by tissue, task, and dataset, with metadata and visualizations. (**C**) *Detail*: comprehensive sample-level metadata, UMAP projections, and statistics. (**D**) *Search*: metadata- and keyword-based retrieval of datasets. (**E**) *Download*: task-, tissue-, and dataset-based data packages with MD5 checksums. (**F**) *Tutorial*: guidance on data formats, labels, and website usage. (**G**) *Analysis*: online tools for cell embedding, cell type annotation, and data mapping.

### Detailed information

Detailed information for each sample selected from the *Browse* or *Search* page is compiled and presented on a dedicated webpage, with each sample ID linked to its corresponding *Detail* page. The *Detail* page encompasses comprehensive metadata for the selected sample, including sample name, dataset, tissue, downstream task, raw data type, organ, sequencing technology, number of cells, number of cell types, number of batches, as well as links to the source data (e.g. NCBI (34), BioStudies (35)) and related publication (Figure 2C). An interactive UMAP projection of cell embeddings generated from EpiAgent is set on the page, enabling users to explore individual cells and toggle labels between cell type, batch, and donor, with associated count distribution displayed for cells, cell types, batches, and donors via pie charts. Additionally, detailed statistics including distributions of fragment counts, non-zero peak counts, and non-zero cCRE counts per cell are presented via histograms.

### Advanced searching

Structured metadata-based filters and keyword-based searches are available at the *Search* page to facilitate flexible queries for sample data (Figure 2D). Users can specify criteria from drop-down menus such as dataset and tissue, and further refine searches by selecting downstream tasks, sequencing technologies, and defined ranges for the numbers of cells, cell types, and batches. A fuzzy search function is also supported, enabling keyword entry to identify potentially relevant samples. Upon submission of a searching query, an interactive metadata table will be presented, enabling direct exploration and download of three types of files for each sample: the cell-by-cCRE matrix, fragment files, and the cell-by-peak matrix. Users can also click on the sample ID in the table to access the detailed web page. In addition, users can click the *Example* button to obtain an exemplary search entry along with corresponding results. Alternatively, a *Download* button located at the bottom right corner of the table allows users to download a.csv file containing both basic dataset information and download links for datasets within the search results, thereby supporting batch data retrieval via command-line tools.

### Data download

In addition to downloading sample files via the tables on the *Search* and *Browse* pages individually, we have developed the *Download* page to facilitate convenient access to prepackaged data (Figure 2E). Specifically, for each task, tissue, and dataset, three file types (cell-by-peak matrix, cell-by-cCRE matrix, fragment file) are archived and compressed into downloadable packages. To ensure data integrity, corresponding MD5 checksum files are provided alongside the downloadable files. We also provide guidance on the usage of various data formats and labeling conventions on the *Tutorial* page, where users can also find a comprehensive tutorial on navigating and utilizing the website (Figure 2F).

Data mapping

### Online analysis and case study

Human-scATAC-Corpus integrates the pretrained EpiAgent model and its supervised derivatives, EpiAgent-B and EpiAgent-NT, into the backend to support three types of online analyses: cell embedding extraction, cell type annotation, and data mapping (Figure 2G). All online analyses require the input data to be provided in.h5ad format as a cell-by-cCRE matrix. To ensure stable performance, the system restricts each individual analysis to a maximum of 10,000 cells, thereby preventing memory overload or excessive computational time for a single task. For larger datasets, users are advised to partition their data into smaller subsets for batch analysis. For each analysis task, users can specify hyperparameters and submit their data; the backend automatically generates a task ID and executes the process. Upon completion, users are notified via the email address they provided and can retrieve the results from the online analysis portal using the assigned task ID. The detailed workflows of the three online analyses and representative case studies are described below.

#### Cell embedding extraction

For this analysis, Human-scATAC-Corpus automatically executes a series of built-in scripts, including data preprocessing, model inference with EpiAgent, and results aggregation (Figure 3A). The generated cell embeddings are stored in the AnnData object under the.obsm field with the key “ cell_embeddings_zero_shot,” and the full AnnData object is returned to the user in.h5ad format. As a case study, we applied this workflow to a randomly downsampled subset of the Buenrostro2018 (30) dataset (1,000 cells), which is also provided as example data on our website. As shown in Figure 3A, our database successfully produced embeddings that capture cellular heterogeneity without requiring local computation, thereby greatly facilitating downstream analyses.

**Figure 3.**
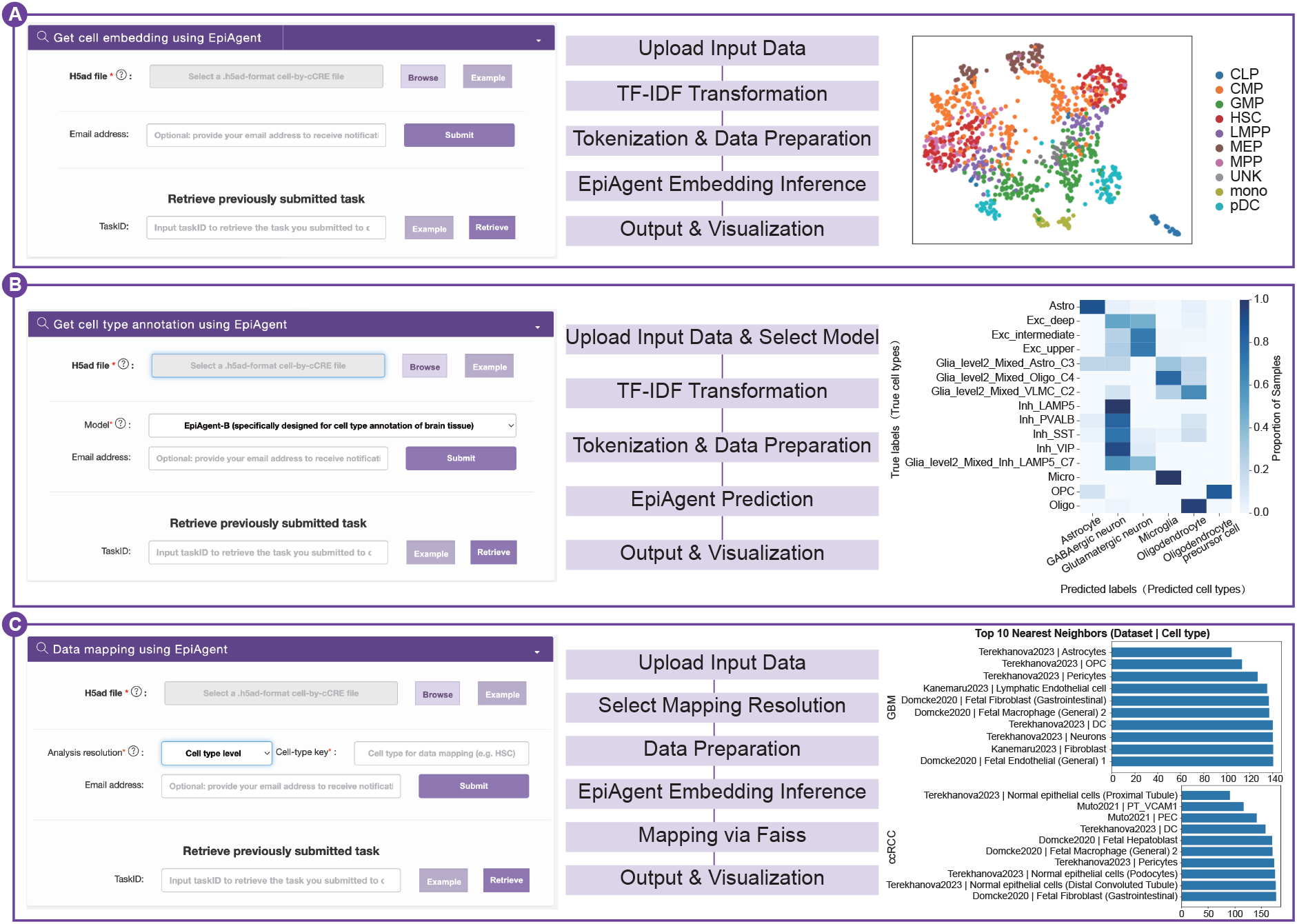
Online analysis in the Human-scATAC-Corpus database. (**A**) Cell embedding extraction: extraction of cell embeddings using the EpiAgent model pretrained on parts of cells in Human-scATAC-Corpus. (**B**) Cell type annotation: supervised prediction of cell types with EpiAgent-B (brain tissue) or EpiAgent-NT (other tissues). (**C**) Data mapping: matching query cells or cell types to normal tissue references within Human-scATAC-Corpus using the EpiAgent model and Faiss library. TF-IDF denotes as Term frequency-Inverse document frequency.

#### Cell type annotation

The pipeline of cell type annotation closely parallels that of cell embedding extraction, with the exception that the backend model is replaced by one of the supervised derivatives, EpiAgent-B or EpiAgent-NT, according to the user’s selection (Figure 3B). Predicted cell type labels are stored in the AnnData object under the.obs field with the key “ predicted_cell_types.” As a case study, we used a randomly downsampled subset of the Li2023 (36) dataset (1,000 cells), also available as example data, and selected EpiAgent-B as the annotation model. As illustrated in Figure 3B, the predicted cell type labels exhibit strong concordance with the ground-truth labels, demonstrating the potential of this function as a powerful tool for preliminary annotation of scATAC-seq data.

#### Data mapping

The data mapping function leverages EpiAgent to associate query cells with reference data, enabling comparisons between disease and normal cells or between in vivo and in vitro cell states, thereby providing insights into pathogenic mechanisms and developmental processes (37). On the backend, reference cells with known type annotations are selected and processed through EpiAgent to obtain their embeddings. For each dataset, the embeddings of cells belonging to the same type are averaged to obtain reference samples.

This function supports two levels of resolution (Figure 3C). In cell-level mapping, each query cell is assigned to the nearest reference sample in embedding space based on Euclidean distance, and the results are returned in a.csv file. In cell type-level mapping, the query cells are first grouped by a user-specified annotation key. For each group, the ten closest reference samples, defined as dataset–cell type pairs, are identified. During the mapping procedure, the Faiss library (38) is employed to accelerate computation. The outputs include an Excel file (with one sheet per group) and distance visualization plots (one per group), all packaged in a compressed archive.

As a case study, we applied cell type-level mapping to a randomly downsampled subset of the Terekhanova2023 (39) dataset (1,000 cells). The results indicated that glioblastoma (GBM) and clear cell renal cell carcinoma (ccRCC) cells were mapped to astrocytes and normal proximal tubule epithelial cells, respectively. These mappings are consistent with the original analyses reported in Terekhanova2023 (39), underscoring the accuracy and reliability of the data mapping function.

## SYSTEM DESIGN AND IMPLEMENTATION

The website is maintained on a Linux-based Nginx web server v1.26.2 (https://nginx.org/) with the backend running PHP (v7.4.33). All metadata are stored in a MySQL database (v8.4.0). We utilize various jQuery and JavaScript plugins including DataTables (v1.10.19), morris.js (v0.5.0), D3.js (v7.8.5), and Chart.js (v3.9.1) to implement and render tables, charts, and visualizations of UMAP projections. The web interface is optimized using the Bootstrap framework (v3.3.7). Besides, the online analysis service powered by the built-in foundation model is currently deployed on a computing node equipped with 4 NVIDIA GeForce RTX 4090 GPUs within a high-performance computing cluster. Considering the lengthy computation time required for the tasks, we designed a parallelization strategy to enable efficient inference across these GPUs. The current Human-scATAC-Corpus website supports most mainstream web browsers, including but not limited to Google Chrome, Firefox, Safari, Microsoft Edge and Opera.

## DISCUSSION

Recent advances in scATAC-seq profiling have produced datasets that are both vast in scale and broad in biological scope, developmental stages (6), spanning diverse tissues (7), and perturbation contexts (9,10,32). This expansion creates the conditions for foundation models to become a central paradigm in scATAC-seq (11,12), yet progress has been constrained by the absence of a curated database that reconciles divergent feature spaces, standardizes metadata, and supports reproducible computation. Recognizing this gap, we developed Human-scATAC-Corpus, a unified ensemble database designed to reconcile these challenges by integrating multiple data representations, spanning broad tissue and perturbation contexts, and embedding a built-in foundation model for online analysis. Our database not only addresses the technical fragmentation derived from characteristics of scATAC-seq data that has hindered integrative analysis, but also establishes an accessible and scalable resource to accelerate the development and application of large-scale foundation models in single-cell epigenomics.

Following the initial release of Human-scATAC-Corpus, we plan to advance the resource along three directions. First, we will extend beyond human data by incorporating data from mouse and other species, establishing a multi-species database. Second, we will systematically curate additional experimental contexts to broaden the range of downstream applications and provide stronger support for method development and benchmarking. Third, we will integrate additional single-cell epigenomic modalities, such as chromatin interactions and DNA methylation, with harmonized metadata and interoperable representations. These expansions are intended to enable large-scale pretraining on comprehensive, multi-omic regulatory signals and to facilitate joint representation learning across species and modalities.

## DATA AVAILABILITY

Users can access all features of Human-scATAC-Corpus without the requirement of registration or login. Users can freely access all data host in Human-scATAC-Corpus at https://health.tsinghua.edu.cn/human-scatac-corpus/.

## AUTHOR CONTRIBUTIONS

R.J. conceived the study and supervised the project. X.C. designed the data storage format of Human-scATAC-Corpus and the frontend the database. X.C., Z.W., and K.L. collected, processed, and annotated the data. Z.G., K.L., Z.W., Q.J., X.C., and Z.L. contributed to the development of the database front-end. X.C., Z.G., Z.W., K.L., and R.J. wrote the manuscript with input from all authors.

## ACKNOWLEDGEMENTS

Not applicable.

## FUNDING

National Key Research and Development Program of China [2023YFF1204802, 2021YFF1200902]; National Natural Science Foundation of China [62273194]; Beijing Natural Science Foundation [L242026].

## CONFLICT OF INTEREST

The authors declare no competing interests.

## Author notes

The authors wish it to be known that, in their opinion, the first four authors should be regarded as Joint First Authors.

## Notes

### Competing Interest Statement

The authors have declared no competing interest.

